# Computational modelling shows evidence in support of both sensory and frontal theories of consciousness

**DOI:** 10.1101/2024.11.02.621640

**Authors:** Kavindu H. Bandara, Elise G. Rowe, Marta I. Garrido

**Affiliations:** Melbourne School of Psychological Sciences, The University of Melbourne, Australia; Graeme Clark Institute for Biomedical Engineering, The University of Melbourne, Australia

**Keywords:** Consciousness, PFC, Computational Modelling, Inattentional Blindness, Dynamic Causal Modelling

## Abstract

The role of the prefrontal cortex (PFC) in consciousness is hotly debated. Frontal theories argue that the PFC is necessary for consciousness, while sensory theories propose that consciousness arises from recurrent activity in the posterior cortex alone, with activity in the PFC resulting from the mere act of reporting. To resolve this dispute, we re-analysed an EEG dataset of 30 participants from a no-report inattentional blindness paradigm where faces are (un)consciously perceived. Dynamic causal modelling was used to estimate the effective connectivity between the key contended brain regions, the prefrontal and the posterior cortices. Then, a second-level parametric empirical Bayesian model was conducted to determine how connectivity was modulated by awareness and task-relevance. While an initial data-driven search could not corroborate neither sensory nor frontal theories of consciousness, a more directed hypothesis-driven analysis revealed strong evidence that both theories could explain the data, with a very slight preference for frontal theories. Specifically, a model with backward connections switched off within the posterior cortex explained awareness better (53%) than a model without backward connections from the PFC to sensory regions. Our findings provide some support for a subtle, yet crucial, contribution of the frontal cortex in consciousness, and highlight the need to revise current theories of consciousness.

## 1.1 Introduction

The necessity of the prefrontal cortex (PFC) in generating consciousness remains hotly debated (Michel & Morales, 2020; Cogitate Consortium, 2025). To date, empirical support has been largely mixed, and the role of the PFC has emerged as a key point of divergence between the current major theories of consciousness. These theories can be split into two theoretical camps suggesting distinct brain regions are involved in conscious perception (Figure 1a and 1b). The sensory (or posterior) family of theories argues that consciousness is accounted for by activity within posterior regions. Frontal theories, on the other hand, propose that the PFC is necessary in the generation of consciousness. While it is beyond the scope of this current paper to compare specific theories of consciousness, we briefly describe them below. For instance, Information Integration Theory, (IIT, Oizumi et al., 2014; Tononi et al., 2016) and Recurrent Processing Theories (RPT, Lamme, 2006; Lamme & Roelfsema, 2000) are two major theories of consciousness that postulates that the posterior regions of the brain are involved in consciousness, although IIT includes some non-sensory posterior regions such as the precuneus and the posterior cingulate cortex (Siclari, 2017). On the other hand, two prominent frontal theories include Global Neuronal Workspace Theory (GNWT [Dehaene & Naccache, 2001; Mashour et al., 2020] and Higher Order Thought Theories, HOTs, [Brown et al., 2019; Lau & Rosenthal, 2011]). While these theories implicate the PFC in consciousness, they do slightly differ in what the precise role of the PFC is in consciousness, with GNWT proposing that the PFC is part of a large, cross-cortical hub, while HOTs (broadly speaking) predicts that the PFC (meta)represent first-order sensory representations.

**Figure 1.**
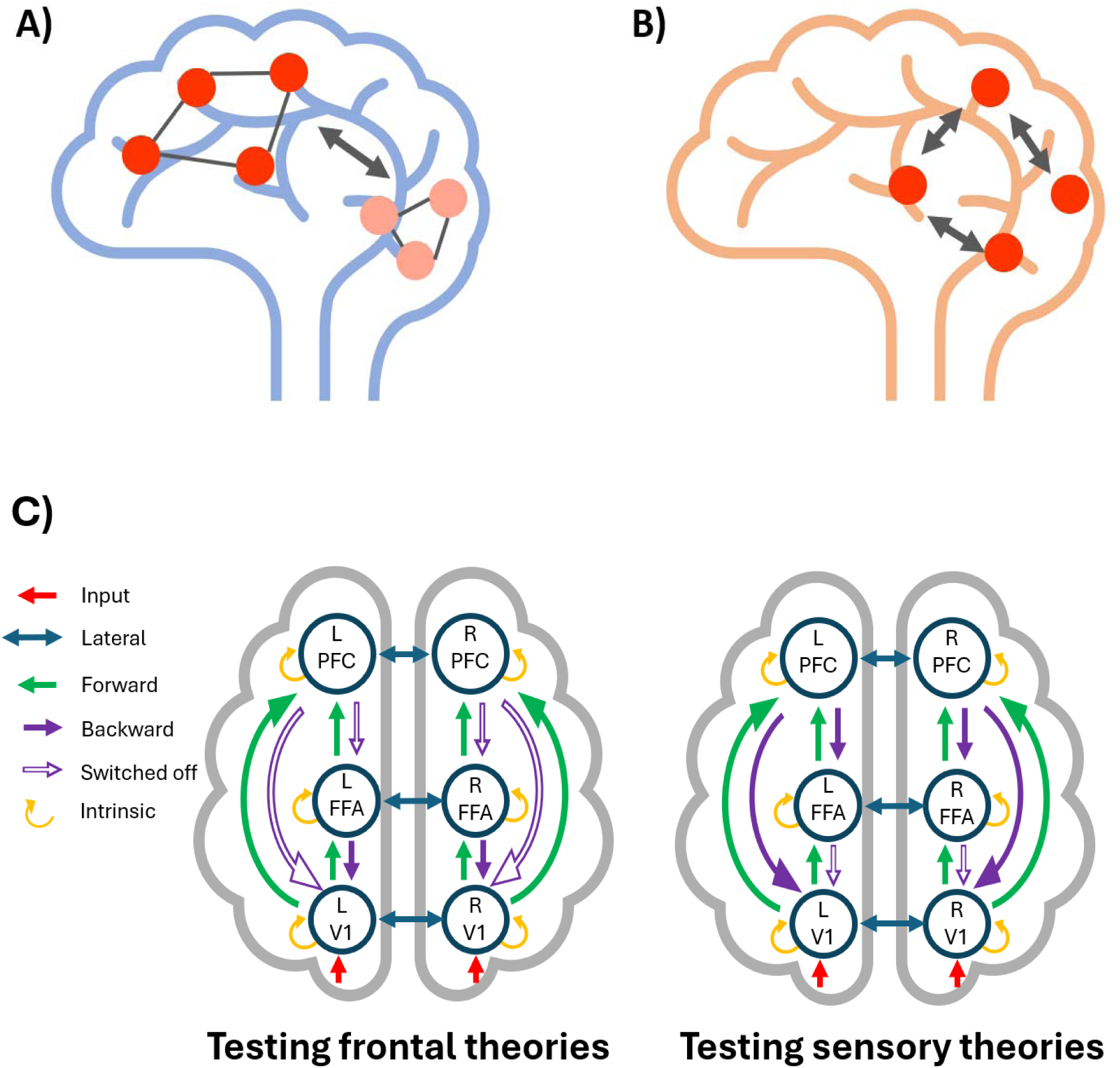
Comparison of Global Neuronal Workspace Theory and Recurrency Theory and the 2 Models Used in our Hypothesis-driven Analysis to Test Predictions of Sensory and Frontal Theories. A comparison of two major theories of consciousness with differing views on the necessity of the PFC in consciousness. A) Global neuronal workspace theory proposes that a frontoparietal network is involved in generating consciousness. When activated, this network broadcasts information across the cortex, enabling access of information from local cortical units. The theory itself predicts more local processors are involved, but for the sake of illustration, a simplified model is depicted here. The key point is that the activation of prefrontal areas is causally involved in the generation of consciousness. B) In recurrent processing theory, feedback activity in recurrent loops within sensory cortices results in consciousness. C) The two models used in a direct model comparison in this paper using two reduced (or nested) parametric empirical Bayesian (PEB) models. These models differed such that key backward connections from either model were fixed at a prior mean of 0 (effectively, switching off the connection). Connections switched off are denoted by hollow arrows in the diagram. On the left model, backward connections from the PFC to lower sensory regions (i.e. PFC-FFA and PFC-V1) are switched off. If losing these connections greatly impacts the model fit, then this suggests that PFC connections are important for awareness, corroborating frontal theories. On the right model, backward connections from the FFA to V1 were turned off. Here, if the model suffers, it suggests that these FFA to V1 connections are important, favouring sensory theories. Here, PFC refers to the prefrontal cortex (Brodmann area 45), FFA is the fusiform face area, and V1 is the primary visual cortex.

Given the well-established role of the PFC in executive control, working memory and other higher-cognitive functions, posterior theories argue that PFC activity is elicited only in accessing and reporting on consciousness, rather than in generating consciousness itself. As the confound of report is difficult to control for, according to sensory theories PFC activity contaminates the true neural correlates of consciousness (NCC; Brascamp et al., 2015; Frässle et al., 2014; Koch et al., 2016; Tsuchiya et al., 2015).

No-report paradigms have been particularly powerful in dissociating the various functions involved in reporting a stimulus from the conscious experience of that stimulus itself. Typically, this is accomplished by delaying the point of report from the time of experience or using other measures which do not rely on explicit report, such as eye-movements (see Frässle et al., 2014). No-report paradigms have already challenged previous interpretations of NCCs; such as the event-related potential component, the P3b, not being a reliable measure of consciousness (Dembski et al., 2021) and the reduction or absence of PFC involvement when stimuli are not perceived (Boly et al., 2017).

Though initial findings from no-report studies challenged the necessity of the PFC in consciousness, recent evidence, particularly when using sensitive multivariate analyses in humans and from fine-grain electrophysiological recordings in non-human primates, adds nuance to this interpretation. There is now accumulating evidence from non-human primates that conscious contents are decodable from the PFC during binocular rivalry (Kapoor et al., 2022; Panagiotaropoulos et al., 2012), and there is partial support for this in humans in a visual masking paradigm (Hatamimajoumerd, 2023; for a recent review see Panagiotaropoulos, 2024). These studies demonstrate that feedforward processing to neuronal populations within the PFC can reliably encode the contents of conscious perception even when post-perceptual confounds related to report are minimized. However, precisely what decoding from the front of the brain means in terms of its role in consciousness is debatable, and as Block (2024) argues, successful decoding from the PFC may not refute nor support frontal theories. It is also useful to note that not all frontal theories characterise the PFC’s role in consciousness as subtle. For example, GNWT predicts a widespread, global ignition of activity from sensory areas and the PFC. What is abundantly clear, however, is that the role of the PFC in consciousness is not well understood and demands further study.

Given the ongoing debate about the role of the PFC in consciousness and the need for no-report paradigms, inattentional blindness (IB) is a valuable paradigm to shed light in this area. IB is a particularly useful paradigm for studying the NCCs underlying conscious awareness as the information entering the visual system is clearly visible and kept constant across participants, yet the IB stimulus only reaches awareness for some people while others fail to detect it (Mack, 2003; for a recent meta-analysis see Hutchinson et al., 2022). Thus, any differences in neural activity between the aware and unaware groups in an IB experiment should, in theory, be required for conscious awareness. What distinguishes IB from other paradigms for probing consciousness is that bottom-up stimulus features are not suppressed in any way; as opposed to masking or manipulating stimulus intensity near perceptual thresholds (e.g., via contrast or duration). In this manner, IB is better able to control for the effects of bottom-up sensory signal strength in consciousness, as the critical stimulus evades conscious perception, not because of stimulus visibility, but because it is unexpected (Hutchinson, 2019).

Given that the IB stimulus is clearly visible, one might expect the corresponding sensory neural signature to be quite strong and potentially propagate to higher-order regions, such as the PFC, even when it is not consciously perceived. Indeed, recent evidence from our lab supports this, showing that feedforward sensory information to the PFC is not modulated by awareness (Rowe et al., 2024). Specifically, using a support vector machine to classify feedforward input patterns to different brain regions, we showed that the classification accuracy for the IB stimulus was above chance across all regions analysed, including prefrontal regions, regardless of participant’s awareness of that very stimulus. This suggests that some baseline level of sensory information makes it to the PFC in IB regardless of awareness. This could be taken as evidence that feedforward processing to the PFC may be (necessary but) insufficient to generate consciousness. However, this does not preclude the possibility that feedback processing from the PFC may be crucial to generating consciousness – as predicted by both GNWT and HOTs.

In this study, we used computational modelling to reanalyse EEG data from an inattentional blindness (IB) no-report task first described by Shafto and Pitts (2015). This task systematically manipulated awareness and task-relevance of faces across three phases. Importantly, all stimuli across all phases were physically identical, with the only manipulation being the task-instructions, allowing us to examine the neural dynamics underlying awareness, while controlling for task-relevance. To adjudicate between frontal and sensory theories, we used Dynamic Causal Modelling (DCM), a technique which estimates the effective connectivity between brain regions, or the directed influence that a brain region has on another (David et al., 2005). This allowed us to investigate the roles of key prefrontal and posterior brain regions, and their interactions, under fluctuating conditions of awareness and task-relevance.

## 2.1 Methods

All data for this study have been kindly provided to us from Shafto and Pitts (2015). For detailed methods please refer to their original paper. However, to provide a brief overview of their experiment (as shown in Figure 2), their task consisted of a three-phase design, where task instructions and expectation were carefully manipulated to isolate task-relevance from awareness. Participants were shown two sets of stimuli – a jittering pattern of lines which, on occasion, would form a face (this was the critical inattentional blindness stimuli), and a set of green circles which formed a ring. Crucially, all trials across phases were kept identical with the only difference across phases being the task-instructions. In phase 1, participants were instructed to ignore the jittering background lines and only respond when one of the dots increased slightly in brightness. After the first phase, participants were given an awareness assessment where they were asked whether they noticed any patterns in the background stimuli. This awareness assessment determined that half of the participants were inattentionally blind (or unaware) of the face and the other half noticed (or were aware) of the face. As it was unclear when participants became aware of the face during phase 1, data from the aware participants were excluded from further analysis. After surprise questioning, in phase 2 expectancy to notice any patterns in the background stimuli increased while participants were instructed to continue to respond to the circle. Another awareness assessment after phase 2 confirmed that all participants became aware of the face in the second phase. This meant that the face remained task-irrelevant, but participants were aware of it. Finally, in phase 3, participants were instructed to respond directly to the faces, and the face became task-relevant.

**Figure 2.**
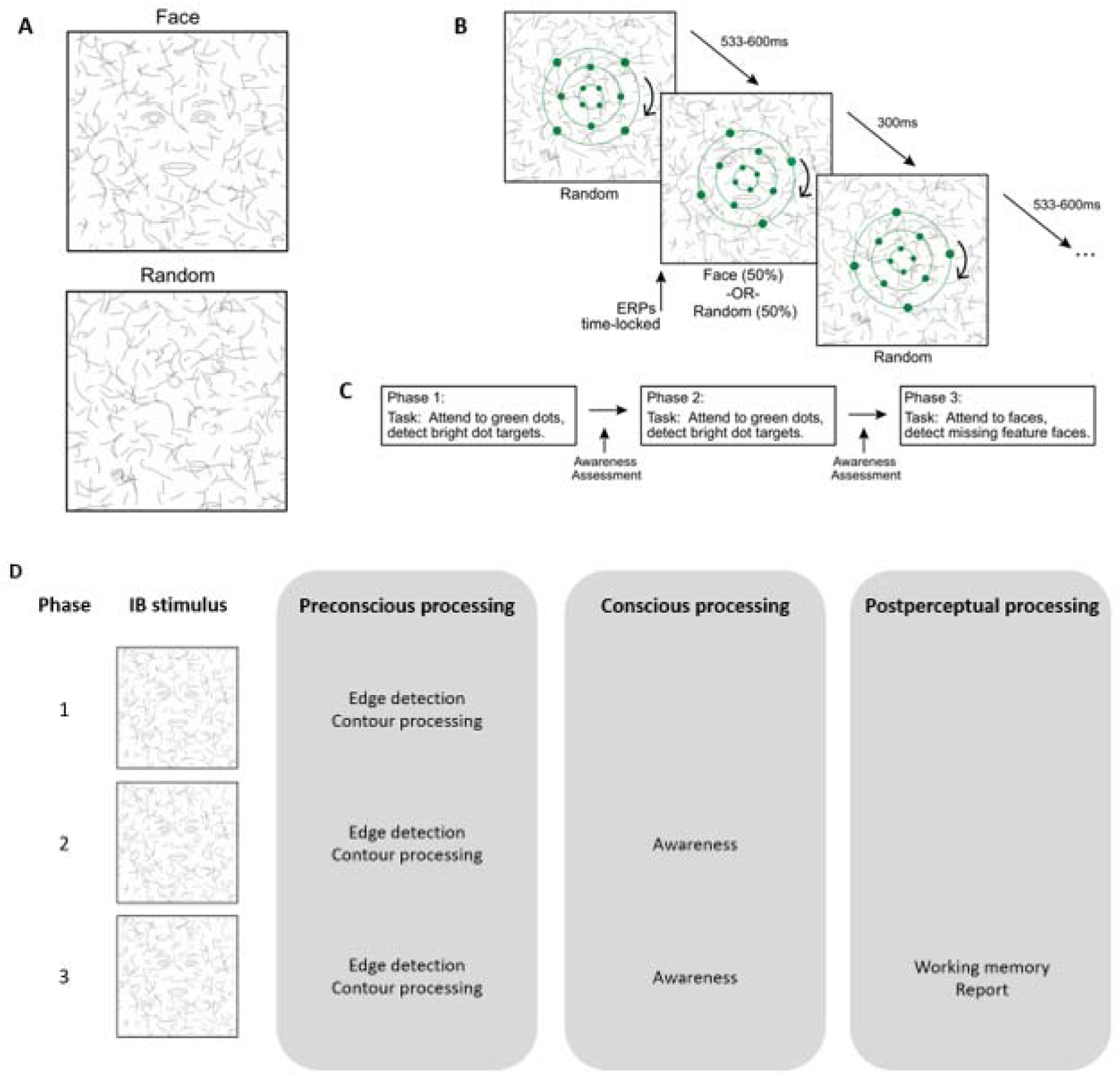
Shafto and Pitts (2015) Inattentional Blindness Task. An overview of the experimental setup in Shafto and Pitts (2015) as well as the key manipulations. A): The two types of background stimuli presented to participants in this task. Here, the face was the critical inattentional blindness stimuli. B): In this task, participants were instructed to respond to an increase in brightness in one of the dots along the green rings. The background lines jittered throughout each phase and participants were told to ignore these jittering lines. On occasion, the jittering lines aligned to form a face – the critical IB stimulus. C): The three-phase design of the task where all trials across phases were kept identical. In phase 1, participants were asked to attend to the green dots, where half the participants were unaware and the other half were aware of the face. In phase 2, all participants became aware of the face but it remained task-irrelevant. Finally, in phase 3, participants were instructed to respond to the faces, and the face became task-relevant. D): Adapted from Pitts et al., 2014, this diagram walks through the different stages of conscious processing throughout the various phases of the task, assuming a participant is inattentionally blind. Preconscious in this sense is used to refer to processing that occurs immediately prior to the conscious representation. This information is supraliminal, in the sense that it *could* be consciously perceived, but it is not made available for report (Dehaene et al., 2006; Pitts et al., 2014).

**Figure 3.**
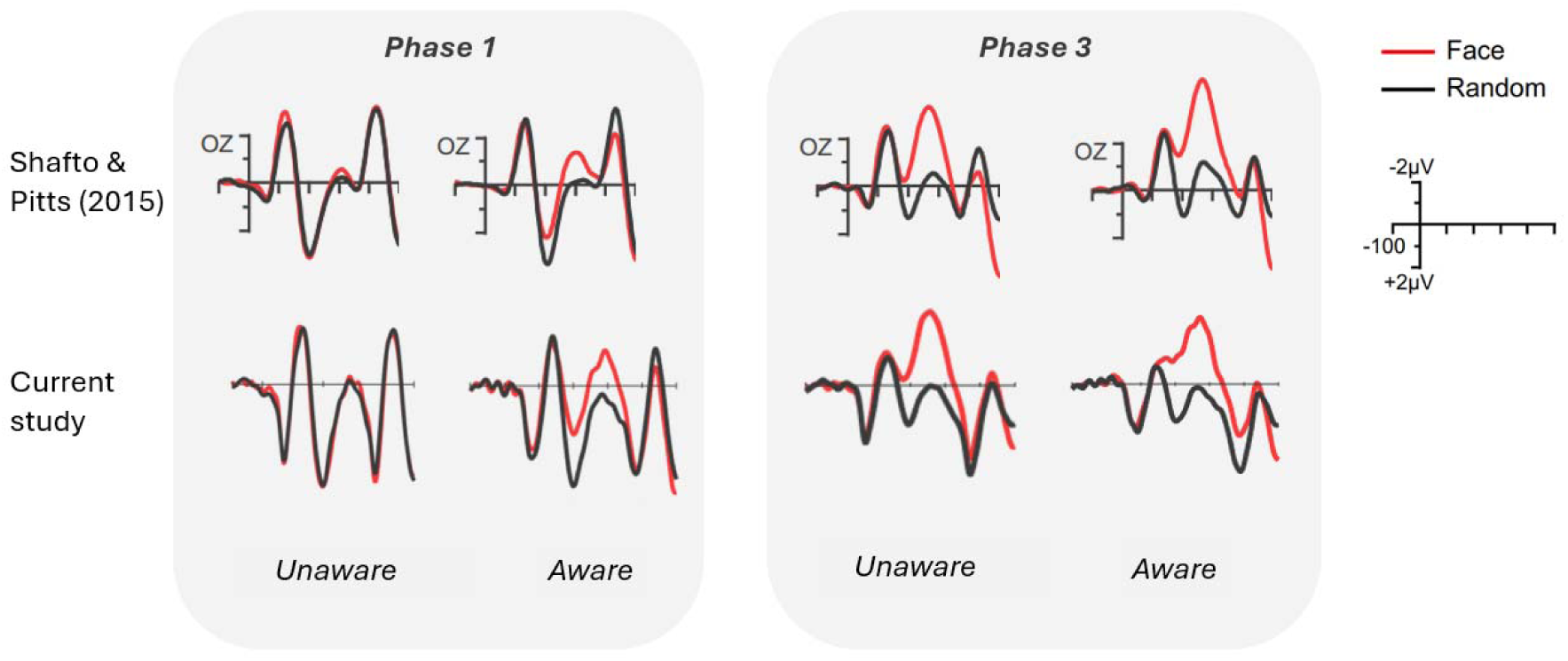
Comparison of Selected ERPs in Current Study and Shafto and Pitts (2015). Comparison of ERPs at electrode Oz in original study (Shafto & Pitts, 2015) with the ERPs computed in the current study after preprocessing. This shows the grand averaged ERPs for unaware and aware participants across phases 1 (when faces are task-irrelevant) and phase 3 (when faces are task-relevant) for face and random trials.

## 2.2 EEG Preprocessing and Analysis

Data preprocessing was slightly different to the original Shafto and Pitts (2015) paper. SPM12 (https://www.fil.ion.ucl.ac.uk/spm/) was primarily used for preprocessing. Data was also preprocessed using a combination of EEGLAB (Delorme & Makeig, 2004; https://sccn.ucsd.edu/eeglab/index.php), and the Chronux software package (Mitra & Bokil, 2007; http://chronux.org/). The raw EEG data originally sampled at 500Hz was first converted to a readable format and then epoched from −100ms to 500ms relative to stimulus onset. Trials containing eye blinks were rejected. These were classified using VEOG channels. Trials containing artefacts, defined as activity greater than 100µV in any channel, and those containing button presses were also removed from analysis. Bad channels were defined as trials which contained these artefacts on more than 20% of trials, and these were also removed from analysis. This led to 16.20% of trials rejected in the unaware group and 11.83% rejected in the aware group. Robust averaging was then performed, and the data was subsequently low-pass filtered at 40 Hz, as robust averaging can introduce high frequency noise. We applied this low pass filter at 40Hz to capture the event-related components known to be evoked by faces. Finally, the data was baseline corrected between –100ms and stimulus onset. Baseline correction is a standard preprocessing step for ERP analysis, including data intended for DCM analysis, to remove slow, non-neural drifts that could otherwise affect the modelling of the evoked response.

## 2.3 Source Reconstruction

To establish the nodes in the DCM models, we can take either a data-driven, or a priori, literature-based approach. As slightly different regions are theorized to be important across different theories of consciousness (even within the same family of theories) a data-driven approach was implemented to guide DCM model specification. Source reconstructed EEG images were used to identify regions of highest activation, which would provide the best chance of finding connectivity changes amongst the regions of interest, should they exist.

To test the hypotheses of frontal and sensory theories of consciousness, at least one PFC node and two sensory ROIs were needed in our network. The need for a PFC node is self-evident; the necessity of the PFC in consciousness is a key point of contention between these two families of theories and the key question of the current study. The two sensory ROIs were provided as a test for recurrent connectivity modulation during awareness, within sensory cortices (a prediction of sensory theories). Sensory nodes were also required as an input point for sensory information to enter the network. These considerations guided the model specification of the DCM analysis.

Source reconstruction on the preprocessed EEG files was conducted in SPM12 (https://www.fil.ion.ucl.ac.uk/spm/), which takes an empirical Bayesian approach to source reconstruction (Phillips et al. 2005). The default SPM template head model was aligned using the fiducial sensors at the nasion, and the left and right periauricular points. This model was first used to run a forward model which computed the effect of source activity on sensor activity. A boundary-element model cortical mesh was used for the forward model, as recommended by the SPM manual (Ashburner et al., 2014). To compute source estimates, this forward model was then inverted using a multiple sparse priors algorithm, optimised using a greedy search routine, which progressively searches for different combinations of source configurations until model evidence no longer improves (Friston et al., 2008). This inversion was conducted using all trials across all conditions and with a time window from – 100ms to 500ms relative to stimulus onset (corresponding to the trial epochs). No specific frequency band was of theoretic interest here, and thus a frequency band window covering the entire frequency range of the data, from 0 to 256 Hz, was chosen. The 3D source reconstructed images were smoothed at a resolution of 12mm^3^ and each image contained 8196 voxels.

The individual source reconstructed data was optimised via a group inversion scheme within SPM. This method takes the individual source estimates and adds an additional constraint to align activity across all participants, but allows the magnitude of activation for sources to vary. This yields better source estimates compared to individual inversion, as it is likely that the same sources are producing activity across all subjects as they are completing the same task (Litvak & Friston, 2008).

## 2.4 Dynamic Causal Modelling

DCM for evoked responses was used first to model how connection strengths modulate in a network when viewing face stimuli (using random stimuli as a baseline) in each subject per phase. The DCM was specified using the nodes obtained from the source-level analysis (see 3.2 Results for details). Here, random and face trials across all phases were inputted as a between-trial effect across a time-window from 1ms to 500ms. An equivalent current dipole model was used to model single dipoles for each source location, as shown in Figure 5. No specific DCM network architecture was specified. Instead, a fully connected network was created, with connections at all levels for forward, backward, intrinsic, and lateral connections. Input connections, however, were restricted to the left and right occipital nodes (i.e., it is well-established that visual information is processed initially in primary visual areas). The modulatory effect of face stimuli was modelled such that all connections were allowed to modulate. These DCMs were inverted per participant per each of the 3 phases (i.e. 3 DCMs per participant to compare phases where faces were either unaware or aware, and task-relevant or irrelevant).

## 2.5 Parametric Empirical Bayes

Parametric empirical Bayesian (PEB) modelling was used to examine how connection parameters in the network were modulated at the group-level across the main effects of awareness and task-relevance. This involved a second-order hierarchical Bayesian model which was conducted over the first-order DCM parameters (Friston et al., 2016). The two main effects of awareness and task-relevance were modelled to observe their corresponding changes in modulatory activity (i.e. the B-matrix in the DCMs) across the network. From here, two types of PEB analyses were performed – an exploratory approach and a hypothesis-driven approach.

### 2.5.1 Exploratory automatic search PEB analysis

First, we took an exploratory approach (also called automatic search), estimating a fully connected model and then reducing redundant connections which did not contribute to model evidence via Bayesian model reduction (BMR; Friston et al., 2016; Zeidman et al., 2019). BMR is an automated procedure that iteratively compares each of the nested models derived from the full model by ‘switching off’ modulatory connections (i.e., setting their prior mean to zero). It then evaluates the evidence for each reduced model. This process effectively searches through a large space of all possible models (including the null model), pruning connections or modulatory effects that do not significantly improve model evidence. The final ‘winning’ model selected by BMR represents the most parsimonious explanation of the data within the explored model space (Friston et al., 2016; Zeidman et al., 2019).

In this step, three regressors were entered into the PEB model. The first regressor modelled the group mean (i.e. face vs. random stimuli), the second modelled how connectivity differed as a function of awareness of the face (0 for unaware and 1 for aware), and finally, the third regressor modelled the difference between task-relevance of the face (0 for task-irrelevant and 1 for task-relevant). For the awareness regressor, phase one was coded as 0 (recall that aware participants phase one data was excluded), while phase two and three was coded as 1. For the task-relevance regressor, phases one and two were coded as 0, and phase three was coded as 1. The full PEB was then estimated and reduced to prune redundant connections.

### 2.5.2 Hypothesis-driven PEB analysis

A second PEB analysis was also performed across a reduced hypothesis space to test specific predictions from either family of theories. Instead of an automatic search, specific parameters within the B-matrix were fixed at a prior mean of 0 (effectively, switching these parameters off). Differences in model evidence when specific connection parameters are switched-off reveal the importance of those connections (Zeidman et al., 2019).

Two candidate models were specified to test frontal and sensory theories, respectively. The interpretation of these models may be counterintuitive at first glance. That is, to determine support for sensory theories, connections from higher to lower sensory regions (VI and FFA) were ‘switched off’ (i.e. fixed at a prior mean of zero), and if this impacted the performance of the model greatly (as measured by model evidence), this would suggest that those connections are important for awareness. On the other hand, to gather evidence for frontal theories, connections from the PFC to both V1 and the FFA were switched off. If such a model suffered greatly by this change in the network architecture, this would suggest that those connections are important for awareness.

As both families of theories agree with respect to how task-relevance would affect activity across the network (i.e. PFC should influence sensory regions), only the awareness main effect was analysed here. The same coding scheme used in the exploratory approach above was used here (i.e. 0 for unaware and 1 for aware). As shown in Figure 6c, two nested PEB models were specified which switched off backward connections from the PFC to sensory regions (testing key connections predicted by frontal theories) or backward connections from higher to lower sensory regions (testing key connections predicted by sensory theories). Model evidence and posterior probabilities of these template PEBs were then compared.

**Figure 4.**
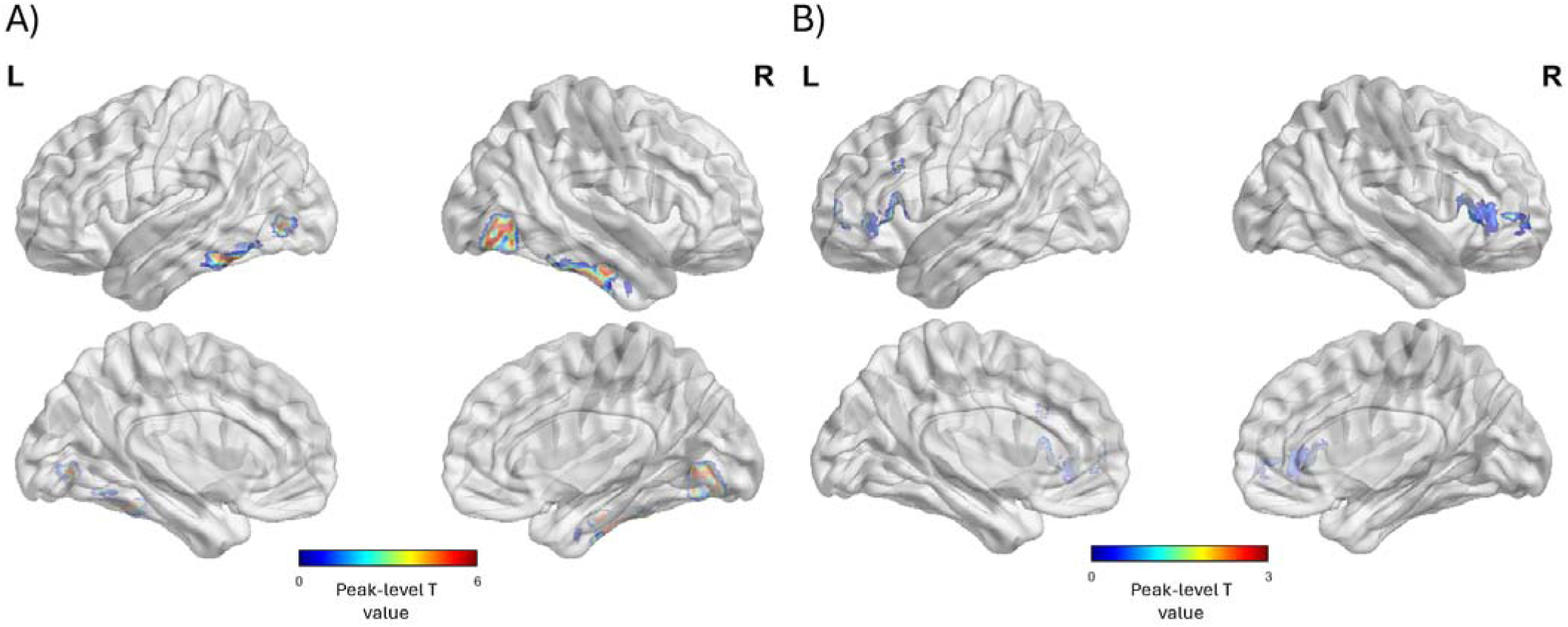
Analysis of Source Reconstructed Images. Significant regions of activation comparing face and random trials in phase three across all participants. A) Source-level contrasts for the interaction between face and random in phase three revealed two clusters across both hemispheres corresponding to occipital and fusiform areas respectively. A family-wise error (FWE) corrected threshold of *p* < 0.05 was used here. B) As no significant prefrontal clusters were found in the first analysis, a less conservative significance threshold of *p* < 0.05 uncorrected was used over an anatomical PFC mask. Peak activity revealed significant clusters at the left and right inferior frontal gyri. L and R denote left and right hemispheres, respectively.

**Figure 5.**
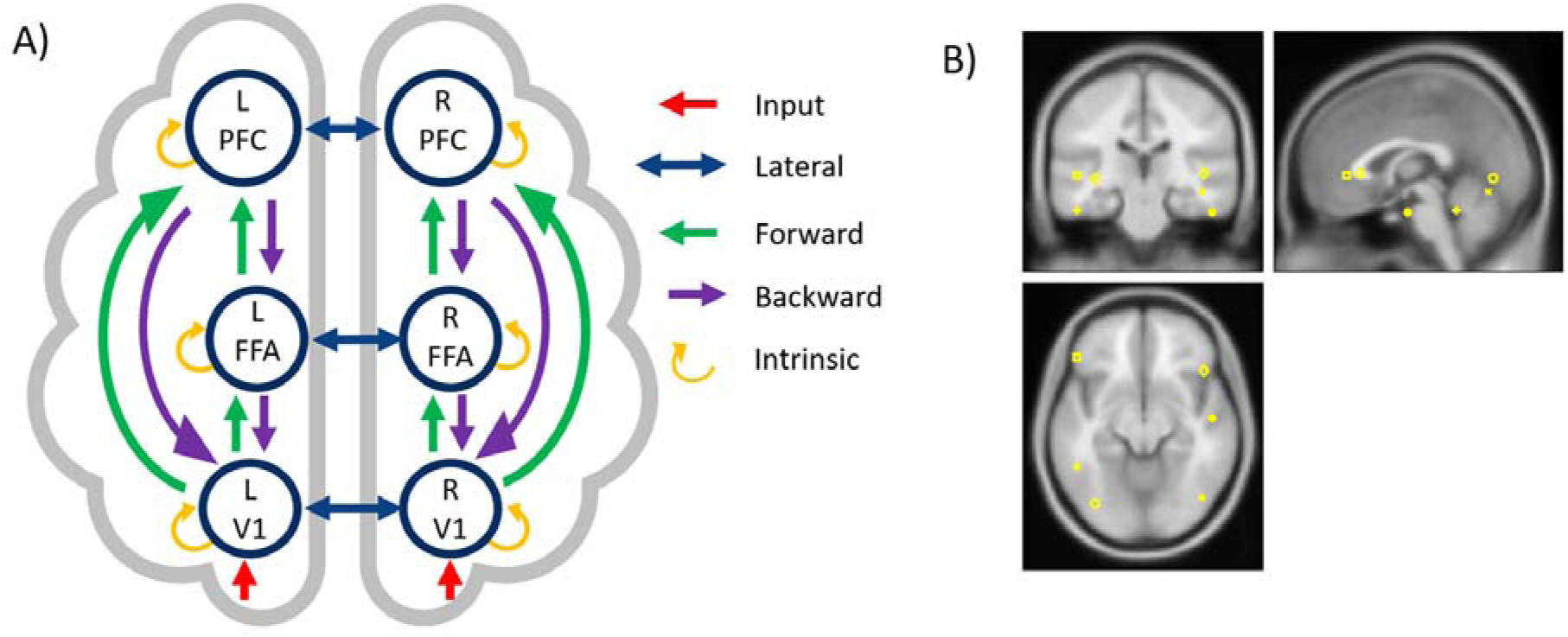
The Full DCM Model Architecture and Dipoles. A) The fully connected DCM architecture used in the DCM analysis. All nodes specified from the source reconstruction image analysis were used and connected at all levels of the architecture. The forward, backward, intrinsic, and lateral connections were free to be modulated by the effect of face. L and R refer to the left and right hemispheres; PFC: the prefrontal cortex node, here, Brodmann area 45 within the inferior frontal gyrus; FFA: fusiform face area; and, Occ: occipital area. B) The dipoles used as nodes in the DCM architecture displayed on a template head model.

**Figure 6.**
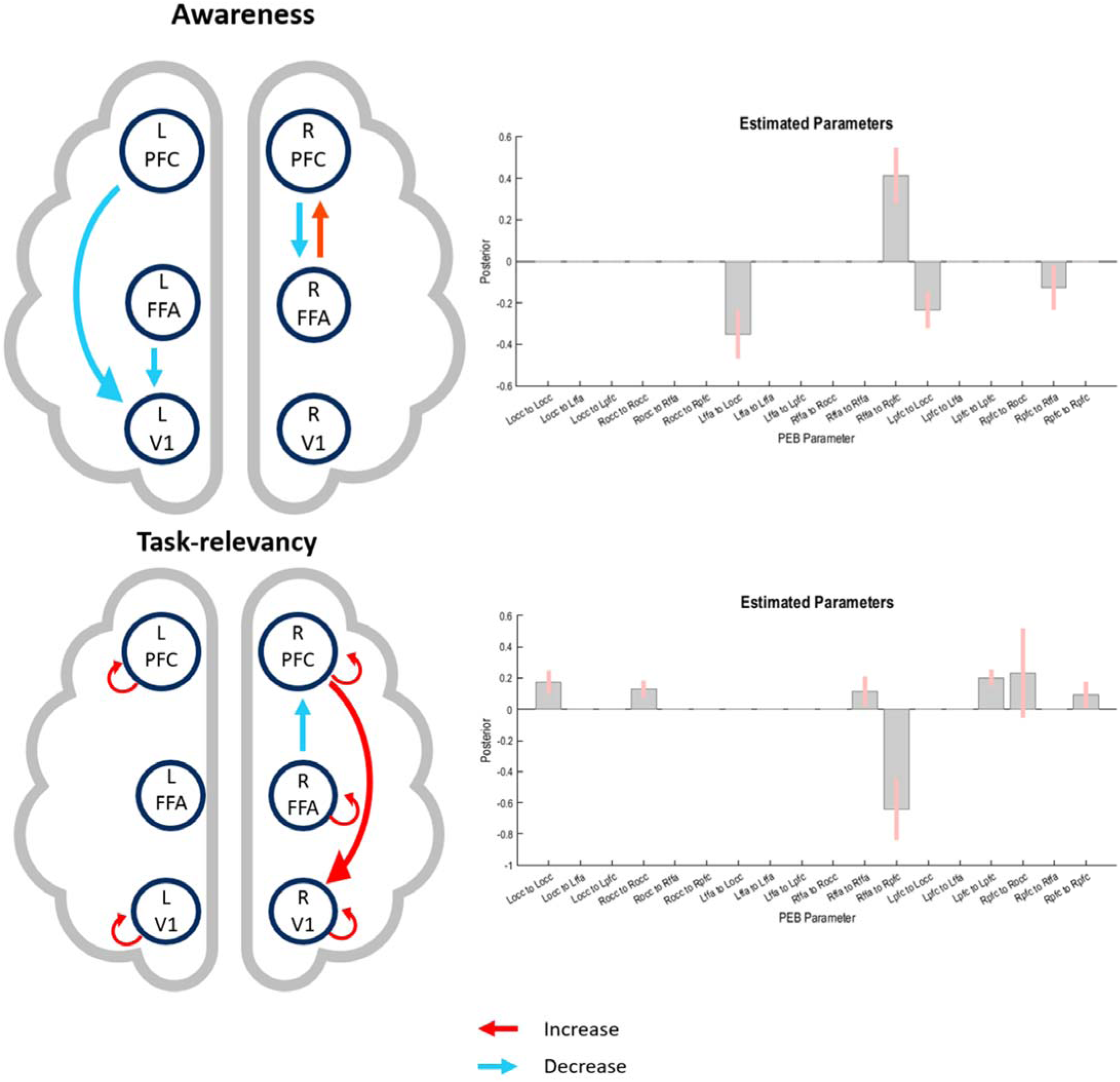
Connection Strengths Across the Network Modulated by Awareness and Task-relevancy using automatic search PEB. This figure shows the results of the winning model via the automated PEB search. The top half shows how the main effect of awareness modulated connectivity across the network. Here, awareness increased connectivity from the right-fusiform to the right-PFC node and decreased from the left-PFC and left-FFA ROIs to the left-occipital node. The bottom half of the figure shows how task-relevance modulated connectivity parameters across the network. Connectivity from the right-fusiform to the right-PFC decreased, while intrinsic connections within the left– and right-occipital, left– and right-PFC all increased (all posterior probability>0.95). PFC refers to the prefrontal cortex (Brodmann area 45), FFA is the fusiform face area, and V1 is the primary visual cortex.

## 3.1 Results

Grand mean ERPs are shown, which through visual inspection, appeared consistent with the ERPs published in the original Shafto & Pitts (2015) paper, despite our slightly different preprocessing. Example ERPs are plotted in Figure 3.

## 3.2 Source-level contrasts

A general linear model was used to determine regions of greatest activation under the experimental conditions. A full factorial model was conducted with a 2 (aware or unaware) x 2 (face or random) x 3 (phase) design. The contrast of interest was the interaction between face and random stimuli in phase three. Phase three was chosen because it is expected that the nodes underlying awareness and task-relevance will exhibit the greatest activity in this condition (where faces are both task-relevant and conscious). Face and random trials were chosen for this comparison to eliminate any common (or baseline) voxel activity. Voxel clusters with the greatest T-statistic were used as nodes in the construction of the DCM. Figure 4 shows the results of these source level contrasts.

Only four clusters with significant activation were found: the right occipital region ([44 –72 –12], peak-level T*_max_* = 6.35, peak-level p*_FWE-corr_* <0.001), right fusiform region ([52 –12 –28], peak-level T*_max_* = 6.2, peak –level p*_FWE-corr_* <0.001), left fusiform region ([50 –48 –26], peak-level T*_max_* = 5.68, peak-level p*_FWE-corr_* <0.001) and left occipital region ([-36 –76 –2], peak-level T*_max_* = 5.66, peak-level p*_FWE-corr_* <0.001). No other significant clusters were found at this adjusted p-value threshold.

In line with our hypotheses set, we identified significant clusters corresponding to one lower-order sensory region and one higher-order sensory-region, from either hemisphere. However, to establish a prefrontal ROI, the same full-factorial model was run but this time with an anatomical mask over the PFC and an uncorrected p-value threshold of *p*<0.05 was used. This PFC mask was created over the Brodmann areas 8, 9, 10, 11, 12, 13, 14, 24, 25, 32, 44, 45, 46, and 47 for left and right hemispheres, respectively (as defined by Fuster, 2015). The most significant prefrontal clusters from these analyses were the left inferior frontal gyrus ([-50 34 0] peak-level T*_max_* = 2.66, peak-level p*_unc_* = 0.004) and the right inferior frontal gyrus ([46 24 2] peak-level T*_max_* = 2.38, peak-level p*_unc_* = 0.009).

It is worth noting that there are individual differences in the reconstructed activity observed in Figure 4. Indeed, individual differences can be captured in the DCM source reconstruction optimisation algorithm when source locations are estimated on an individual-by-individual basis. For these data, however, this approach decreased model evidence (i.e. free energy, results not shown) because the added value in model fit accuracy did not outweigh the penalty of increased model complexity. Hence, we report the results of the best model, which considered source location parameters as fixed effects (not taken to the second level, Zeidman et al., 2019).

## 3.3 DCM and PEB analysis

The PEB analysis revealed connectivity changes across the inverted DCMs as a function of awareness and task-relevance. To reiterate, two approaches were taken here – an exploratory automatic search to determine how connection strengths are modulated by awareness and task-relevance, and a hypothesis-driven search to examine whether a frontal or sensory model best explained the data.

The automatic search revealed how awareness and task-relevance modulated the strength of face-specific connections across the network and throughout the three phases. The estimated group-level connection strengths as a function of awareness and task-relevance is shown in Figure 6 for the winning model. For the main effect of awareness, connectivity increased from right fusiform to the right PFC. Though this activity was lateralised, this may provide partial support in line with the idea of global ignition from PFC to sensory areas postulated by a prominent frontal theory (GNWT; Mashour et al., 2020). We also saw decreased connectivity from the left PFC and left fusiform nodes to the left occipital node as a function of awareness (posterior probability > 0.75). For the main effect of task-relevancy, we found that connectivity from the right fusiform to the right PFC decreased, while intrinsic connections within the left and right occipital, left and right PFC, and right fusiform all increased (posterior probability > 0.75).

As these results did not provide clear support for predictions from either theoretical camp, a more directed hypothesis-driven approach was taken to constrain the model space to test specific hypotheses. When comparing two nested PEB models to explain modulatory changes in awareness, with either FFA to V1 connections switched off (to assess sensory theories) or PFC to both FFA and V1 connections switched off (to assess frontal theories), to, there was a slight preference for the model with FFA to V1 connections switched off with a posterior probability of 53% compared to 47% for the model with PFC backward connections switched off. Hence, slightly favouring the frontal theories, Crucially, to ensure that these backward connections were indeed important in contributing to model evidence, a null model (where all backward connections were switched off) was added into the model space. This null model performed poorly and had a negligible effect on the model probabilities, confirming the importance of these backward connections in the model.

## 4.1 Discussion

This study aimed to investigate the function of the PFC in visual consciousness by investigating how effective connectivity modulates within a network of posterior and frontal regions during IB. We conducted second-order modelling using PEB on first-level DCMs to investigate how particular regions contribute to consciousness under conditions of awareness and unawareness. Through an exploratory search based on Bayesian model averaging, we found inconclusive results which sided with neither frontal nor sensory theories of consciousness, albeit a feedforward connection from the right FFA to PFC provide partial support for a frontal theory (as in GNWT; Mashour et al., 2020). Through a subsequent hypothesis-driven approach, we found evidence supporting both theories of consciousness, with a very slight preference for frontal theories (53% for frontal vs 47% for sensory).

The exploratory automatic search PEB results revealed that in the winning model, awareness of the face increased connectivity from the right fusiform to the right PFC nodes and decreased from the left PFC to the left occipital ROIs. These results do not align with either family of theories. Various sensory theories predict feedback connectivity would increase under awareness within higher and lower-sensory regions – however this was not observed. Secondly, though frontal theories predict the PFC would increase feedback connectivity to sensory areas, instead a decrease was observed. Having said that, an increase in connectivity was observed from the right fusiform to the right PFC, providing partial support for the idea of global ignition via the PFC as predicted by GNWT (Mashour et al., 2020), one of the frontal theories.

The results for task-relevancy were similarly unexpected. For the main effect of task-relevancy, we observed a decrease in connection strengths from the right fusiform to the right PFC, while intrinsic connection strengths for the right occipital, left PFC, and right PFC all increased. This was unexpected as both theories predict PFC connectivity to sensory regions should increase with task-relevancy.

As these results did not clearly align with predictions from either theoretical camp, a more directed hypothesis-driven approach was taken to limit the model space to test specific hypotheses. Two candidate models were specified to test frontal and sensory theories, key connections were switched-off, and model evidences was computed and compared. The results from this analysis revealed that both sensory and frontal feedback connections are necessary for consciousness, with a very slight preference for frontal theories, as the model which did not allow backward connections from the PFC to modulate, had a relatively lower posterior probability.

At face value, these findings provide some support for frontal theories, albeit only slightly. Our results reveal that a model with key connections from the PFC to higher– and lower-sensory regions switched off, fare slightly worse compared to a model where only higher-to lower-sensory connections are switched off. This suggests the backward connections from the PFC may be important for awareness and the model suffers without them. This could suggest a subtle influence of the PFC in consciousness as predicted by frontal theories.

Thus far, we have discussed frontal theories as a family of theories; however, these theories do differ in how frontal areas precisely instantiate consciousness. Most notably, various proponents of higher-order thought theories advocate for a relatively subtle, yet essential, role of the PFC in consciousness (Michel & Morales, 2020; Morales & Lau, 2020; Odegaard et al., 2017). Higher-order thought theories propose that consciousness arises when higher-order regions (meta-) represent lower-order perceptual. These theories hypothesise that the neural architecture involved in targeting first-order representations may not be that extensive, and quite sparse (Lau, 2019). Accordingly, various findings showing no or little PFC activity under conditions of (non-reported) awareness are argued to be due to traditional imaging methods (e.g., fMRI, MEG) lacking the sensitivity to detect this subtle activity (Michel & Morales, 2020; Morales & Lau, 2020).

There are several studies which support the view that activity in the PFC is reduced, but not absent, during awareness in no-report tasks (Dellert et al. 2019; Frässle et al., 2014; Hatamimajoumerd et al., 2022; and using more sensitive measures in Noy et al., 2015; and Fang et al., 2023). Using a visual masking paradigm, a recent study by Hatamimajoumerd et al. (2022) provided partial support for this view. They demonstrated that fMRI activity from frontal regions was modulated by the requirement to report on a visible stimulus; such that, frontal activation was elicited in report conditions but was absent in no-report conditions.

However, when using more sensitive analysis techniques (i.e. multivoxel pattern analysis), a classifier could reliably decode conscious awareness from frontal regions, irrespective of the requirement to report. This could be suggestive of a subtle, albeit crucial, role of the PFC in consciousness, as predicted by some frontal theories. Nonetheless, this conclusion should be treated with caution as frontal areas were the only region (out of the regions studied) where decoding accuracy was significantly modulated by report. More recently also, the adversarial collaboration by the Cogitate Consortium (2025) found mixed results for the role of the PFC even within the same dataset, as decoding performance in the PFC depended on the stimulus feature being analysed. Taken together it may be that, under no-report conditions, awareness is indeed decodable, but to a lesser extent, due to a baseline level of executive processing involved in completing the task (Block, 2019).

The current study adds to the mixed results from univariate activation comparisons and varying interpretations from decoding analyses. One possible explanation for a lack of PFC activity may be that content-specific PFC activity might be subtle, sparsely distributed, or occur at a finer spatiotemporal scale than readily captured by macroscale methods like fMRI or scalp EEG (Panagiotaropoulos, 2024). However, precisely how this would be empirically instantiated requires further refinement and technological advancement. Setting aside the limitations of current neuroimaging techniques, the so-called ‘pointer’ (or relational) frontal theories of consciousness may provide an alternative explanation. These theories posit that the PFC simply acts as a router (or points) to perceptual contents in sensory areas and hence, these versions of frontal theories do not predict that decoding of perceptual contents from the PFC is required in the first place (Block, 2024; Michel, 2022). In these interpretations, whether perceptual contents are decodable from the PFC is beside the point, as the true role of the PFC in consciousness may be, for example, to send a signal when a posterior belief is updated about what an object is. However, given that our current study was designed to investigate network interactions for seen and unseen faces (hypothesising enhanced feedback connectivity from relevant areas during seen conditions), and yet still our results failed to provide clear differentiating evidence in explaining the PFC’s precise role, perhaps underscores the need for further theoretical and empirical refinement. Therefore, to conclusively move toward resolving the ambiguity around the exact role of the PFC in consciousness (whether that be subtle representation, a relational signalling function, or another mechanism altogether), necessitates further study employing more sensitive measurement techniques and more refined computational modelling approaches tailored to test the specific models predicted by these distinct theories.

Our results, particularly the hypothesis-driven comparison, suggest feedback connections from both the PFC and higher-sensory regions may play a role in consciousness. Given that our results show support for both theoretical camps, it may be fruitful to consider precisely what different theories of consciousness target as their explananda and how their distinct goals might account for our results (Storm et al., 2024). Specifically, consciousness has often been conceptually distinguished into phenomenal consciousness (the subjective quality of experience) and access consciousness (the availability of that information, e.g. for report; Block, 1995). Theories emphasising frontal involvement often conceptualise consciousness in terms of its access, whereas sensory theories tend to disentangle the two and claim phenomenal consciousness primarily occurs in posterior regions. In paradigms like the one used here, where stimuli become consciously perceived (e.g., across Phases 1 and 2), participants likely gain both phenomenal experience and cognitive access simultaneously.

This overlap makes it challenging to isolate whether the observed neural activity from frontal and/or sensory regions reflects one type of consciousness, or the other, as it may be the case that both types of consciousness are co-occurring and hence, emerge in our findings to support both theories. To resolve this ambiguity, future experiments could utilize paradigms that may offer the ability to empirically dissociate phenomenal and access consciousness (Hutchinson et al., 2022; Amir et al., 2023). This would enable a targeted test of whether frontal feedback distinctly tracks cognitive access, as predicted by some theories.

We also note that the relatively weaker reconstructed group source activity in the PFC compared to sensory areas, could have been due to the known functional heterogeneity and mixed selectivity within the PFC. Although PFC engagement was observed (albeit with a liberal threshold), the spatial topography of frontal activation within our participants was somewhat varied. To determine whether the weaker statistical effect in PFC was due to weak activity, greater topographical variability across individuals, or both, we inspected both the activity maps and their overlap across participants (see Supplementary Figures 1 and 2).

Indeed, the reconstructed activity in the PFC was weaker in sensory areas. A consistent pattern of activation was found in Dellert et al. (2021) using a similar IB task, but with simultaneous EEG-fMRI. That is, they noted strong occipital and fusiform activity but comparatively weak (or absent) frontal activation in their group-level statistics. They hypothesise weaker frontal activation may have been due to interindividual heterogeneity in their sample. Only a small cluster in the right inferior frontal junction emerged in the group-level analysis. Notably, this region lies adjacent to the inferior frontal gyri observed in our source-level contrasts. This shows using different imaging modalities, samples, and analyses, both our study and Dellert and colleagues’ study converge on a similar pattern of activity; i.e. the same frontal locus emerges at the group-level, but its activation is comparatively weak.

This may well explain why our attempt to account for individual variability by loosening prior variance at the sources in our first-level DCMs analysis reduced overall model evidence. This is likely because the increase in the number of parameters (one PFC source location per individual, rather than one overall for the group) was penalised more so than their added benefit in model fit.

In sum, our findings broadly show support for both sensory and frontal theories of consciousness. Our approach is timely and coincides with a broader push in the field toward pitting theories of consciousness against each other in a systematic and unbiased way. By directly comparing key points of divergence between theories, this approach hopes to accelerate progress toward an understanding of consciousness (Melloni et al., 2021). This has been best exemplified by a recent large-scale adversarial collaboration (Cogitate Consortium, 2025). Additionally, as DCM and PEB are Bayesian approaches, our results can be used within the framework outlined by Corcoran et al. (2023) to formally evaluate the relative support this dataset provides for competing theories of consciousness, using the standardized Bayesian metric of log model evidence. Importantly, given the (relatively) inconclusive results of the recent adversarial collaboration, ours, and that of Rowe et al. (2024), we argue for a need to revise the current theories of consciousness, including re-examining precisely what the role of the PFC is in consciousness.

## Acknowledgements

The authors would like to thank Dr. Peter Zeidman and Dr. Kit Melissa Larsen for their extremely helpful guidance during the data analysis process, specifically in relation to the DCM-PEB analysis. We would also like to thank Dr. Juliet Shafto and Dr. Michael Pitts for collecting the data and sharing this data with us for reanalysis. KB was supported by the Melbourne School of Psychological Sciences PhD Studentship and the Australian Government Research Training Program Scholarship.

## Data Availability

The data underlying this article will be shared on reasonable request to the corresponding author.

