## Supplementary figures for "Computational modelling shows evidence in support of both sensory and frontal theories of consciousness"

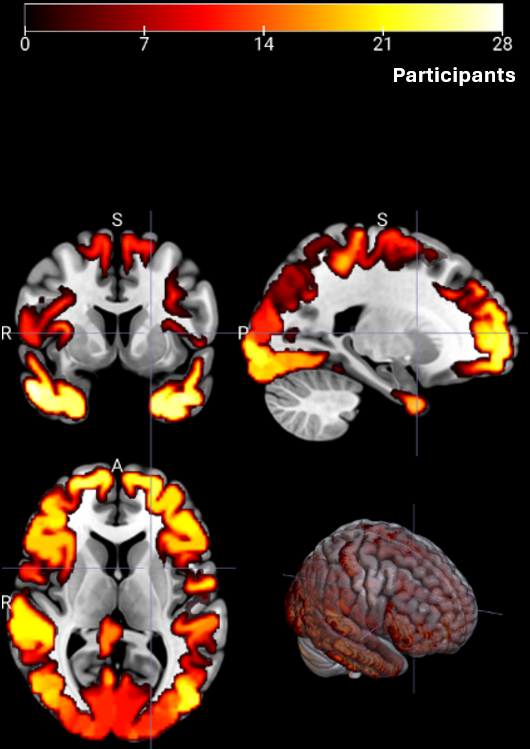

**Supplementary Figure 1**. *Overlapping of Binarised Source Reconstructed Activity Across Participants for Faces vs. Random.* This figure shows the number of participants with any amount of activity in a given brain voxel when comparing faces vs random images from our source reconstruction. Specifically, more yellow/white means more participants had some activity in that region.

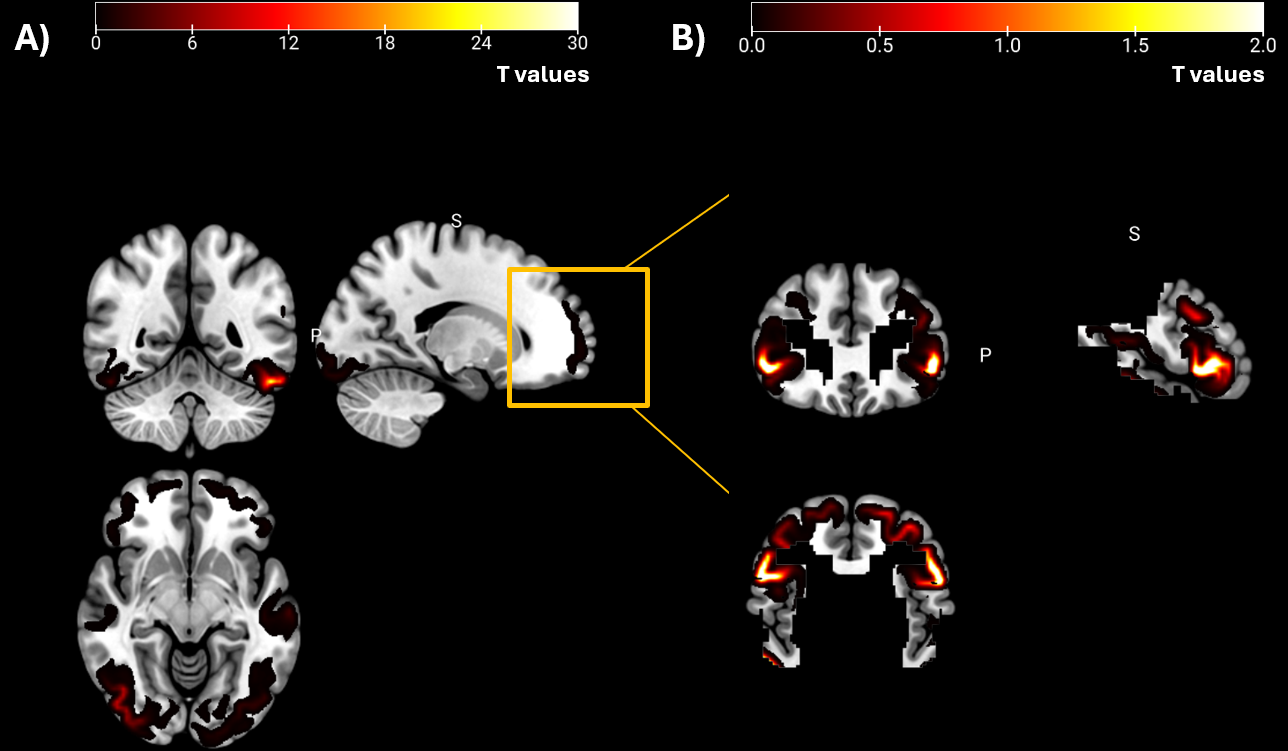

**Supplementary Figure 2**. *Averaged Difference of Source Reconstructed Activity Across Participants for Faces vs Random* This figure shows the average of source reconstructed activity when comparing faces and random trials. A) shows the average activity across the entire cortex, revealing stronger activity in occipital and fusiform areas. B) shows the same contrast but only in a masked region of the PFC. Note the difference in the maximum range of the legend reveals that activity in the PFC was weaker than in posterior regions.
